# Genome-wide analysis reveals signatures of selection for gait traits in Yili horse

**DOI:** 10.1101/471797

**Authors:** Ling-Ling Liu, Chao Fang, Jun Meng, Johann Detilleux, Wu-Jun Liu, Xin-Kui Yao

## Abstract

High quality gaits play an important role in many breeding programs, but the genes of gait were limited at the moment. Here, we present an analysis of genomic selection signatures in 53 individuals from two breeds, using genotype data from the Affymetrix Equine 670K SNP genotyping array. The 11 selection regions of Yili horse were identified using an *F*_ST_ statistic and XP-EHH calculated in 200-kb windows across the genome. In total, 50 genes could be found in the 11 regions, and two candidate genes related to locomotory behavior (*CLN6, FZD4*). The genome of Yili horse and Russian horse were shaped by natural and artificial selection. Our results suggest that gait trait of Yili horse may related to two genes. This is the first time when whole genome array data is utilized to study genomic regions affecting gait in Yili horse breed.

## Introduction

Artificial selection in horse has resulted in divergent breeds that are specialized for either sport or milk production or raised as dual-purpose breeds. Such selection strategies are likely to have imposed selection pressures on particular regions of the genome that control these traits. Such as the Arab horse was used to endurance races (Cervantes *et al*. 2009).

Gait is an important factor that affecting the pattern of horse locomotion. A few Chinese horse breeds have a natural talent for gaits other than the common walk, trot and gallop, such as pace, which is a lateral two-beat gait (Hou 2012). Yili horse was called “tianma” and is a breeding breed in Xinjiang. This breed emerged about 2000 years ago in Western Han dynasty. In 1956, it was officially called “Yili horse” from the crossing of Thoroughbred, Don horse with Kazakh horse of native horses selected by local breeders (Yang *et al*. 1987). Pace is a two-beat gait in which the horse moves the two legs on the same side of the body in a synchronized, lateral movement in contrast to the trot, where the diagonal front and hind legs move forward and backward together (Andersson *et al*. 2012). In horses used for harness racing, breeders selected for the ability of a horse to trot or pace at high speed.

In recent years, methods for detecting selection signals using chip data have been widely used in livestock (Liu *et al*. 2018; Zhao *et al*. 2015; Ai *et al*. 2014; Gutiérrez *et al*. 2017). *F*_ST_ was used to detect the degree of differentiation between populations by the allele frequency (Wright 1950). XP-EHH was developed by Sabeti *et al*. on the basis of extended Haplotype Homozogyity (EHH) (Sabeti *et al*. 2007), which is efficiently to detect selected site. The companion study shows that selective signal method that based on haplotype-frequency was more accurate and reliable than method which based on single-site (Chen 2016). These two methods were widely used in domesticated animals, as in the cases of sheep (Yuan 2016; Zhu 2017), horse (Gu *et al*. 2009). However, the single *F*_ST_ method may generate bias and false positives. Two methods would be more reliable if a reference population with a similar demographic history is available and if the allele under selection is close to fixation in one of the populations (Wagh *et al*. 2012). To minimize the bias and false positives, we combined these two approaches for the detection of selection. Genes related to gait traits are less reported, only the DMRT3 gene is described as the main gene involved in the determination of gait phenotypes in different horse breeds (Niina 2013).

The aim of this study was to identify selected regions related to gait trait of Xinjiang Yili horse breed based on Affymetrix Equine 670K high-density chip using *F*_ST_ and XP-EHH.

## Materials and Methods

### Animals

A total of 53 horses from two populations were used in this study, including 40 Yili (YL) horse and 13 Russian (RUL) horse, all from Zhaosu county of Yili state of Xinjiang (East longitude 81°08’, latitude 43°15’). Jugular blood of 5mL was collected from each animal in tubes treated by ACD anti-coagulation and stored at −20 °C. The specimen was released.

### Quality control of SNPs

DNA extraction and SNP genotyping were performed at the CapitalBio Technology in Beijing. The Affymetrix equine SNP670 chip was used to genotype 53 horses. The SNP data was filtered by PLINK (v 1.9) software (Purcell *et al*. 2007). The following criteria were applied to all SNPs: (1) SNPs with maximum missing rate < 0.99 were removed; (2) SNPs were filtered based on a minor allele frequency (MAF) < 0.03. (3) SNPs were filtered when there was no known autosomal genomic location, (4) SNPs were filtered when the call rate < 0.9, (5) SNPs were filtered that in Hardy-Weinberg equilibrium (*P* < 10^-3^).

Beagle (v 4.0) was used to impute missing genotypes and infer the haplotype phase (Browning *et al*. 2007).

### Population structure

Principal component analysis (PCA) was performed using PLINK1.9 and the ggplot2 package in R (v3.4.4) was used to draw PCA figure (Wickham 2015). We removed SNPs in linkage disequilibrium in PLINK 1.9 with the command (--indep-pairwise 50 5 0.2).

### *F*_ST_ test

*F*_ST_ was used to measure the distribution of genetic diversity among populations (Wright 1950; Weir *et al*. 1984), which uses the variance analysis to get unbiased estimates of Wright’s *F*_ST_. *F*_ST_ between RUL and YL breeds in 25-kb windows with sliding and each site were calculated by Vcftools (v0.1.15) (Danecek *et al*. 2011). We selected the top 0.5% of SNPs (*F*_ST_>0.1699) as candidate sites.

### XP-EHH test

For estimating XP-EHH, we use selscan (v1.1.0b) to confirm the selected sites detected by *F*_ST_ between RUL and YL populations (Szpiech *et al*. 2014; Pickrell *et al*. 2009), which is a more powerful method for selective sweeps that have above 80% frequency (Voight *et al*. 2006).

The top 0.5% of SNPs (XP-EHH > |-1.9405|) were selected as “candidate sites”. RUL horse as the experimental population and YL horse as the control population.

### Bioinformatics analyses

The 200 kb upstream and downstream of candidate sites were defined as the boundary of candidate regions. Horse annotated genes were searched via BioMart database of the ensemble genome browser (Arek 2011).

GO terms, including biological process, cellular component, and molecular function, and KEGG pathway enrichment analyses were performed on the genes of identification of selected candidate regions, using the function annotation clustering tool from the Database for Annotation, Visualization and Integrated Discovery (DAVID v6.8, http://david.abcc.ncifcrf.gov/) (Huang *et al*. 2009). Corrections for multiple testing were performed by applying the Benjamini-Hochberg method (Benjamini *et al*. 1995), and GO terms and the KEGG pathways were considered signifcant at a P value lower than 0.05.

Venny2.1.0 (http://bioinfogp.cnb.csic.es/tools/venny/index.html) was used to draw a Venn diagram of overlap sites of two methods.

## Results

### Genotype and quality control results

After quality control, 143,652 SNPs were removed due to missing genotype data (--geno), and 124,447 SNPs removed due to MAF, 23,702 SNPs on Un and X chromosome, 1,700 SNPs removed, which were in Hardy-Weinberg equilibrium (*P* < 10^-3^). The remaining 377,295 SNPs were used for subsequent analysis.

### Population stratification assessment

Results of the PCA are shown in Figure 1. PC1 explained 3.13% of variance, and PC2 explained 1.93% of variance. Result showed that RUL and YL horse were separated by PC1 and there was no mixed in two populations, which provided foundation for the subsequent selection of signal detection.

**Figure 1.**
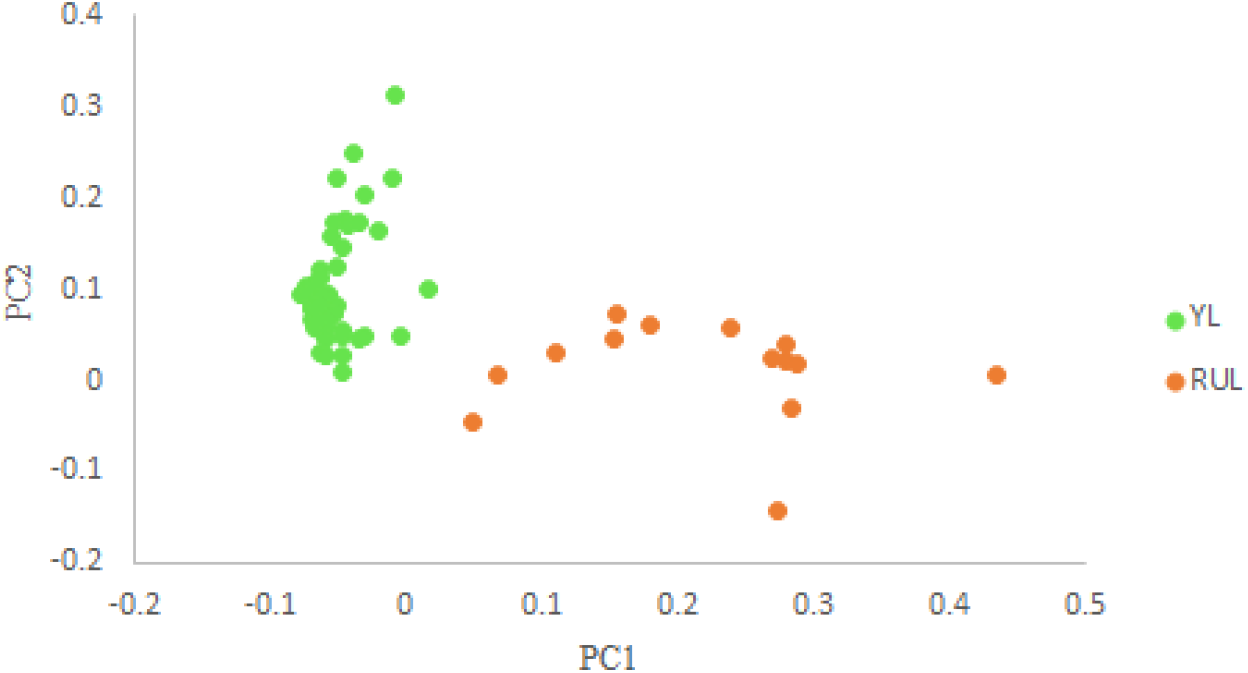
Principal component analysis (PCA) of RUL and YL horse breeds

### *F*_ST_ test

Vcftools was used to calculate *F*_ST_ of each SNPs between two populations. The *F*_ST_ value (*F*_ST_ = 0.1699) was set as the threshold, and the highest *F*_ST_ of SNP (AX-103963716, *F*_ST_ = 0.56) was located on chromosome 1. A total of 1,886 sites were selected. Manhattan map was showed in Figure 2.

**Figure 2.**
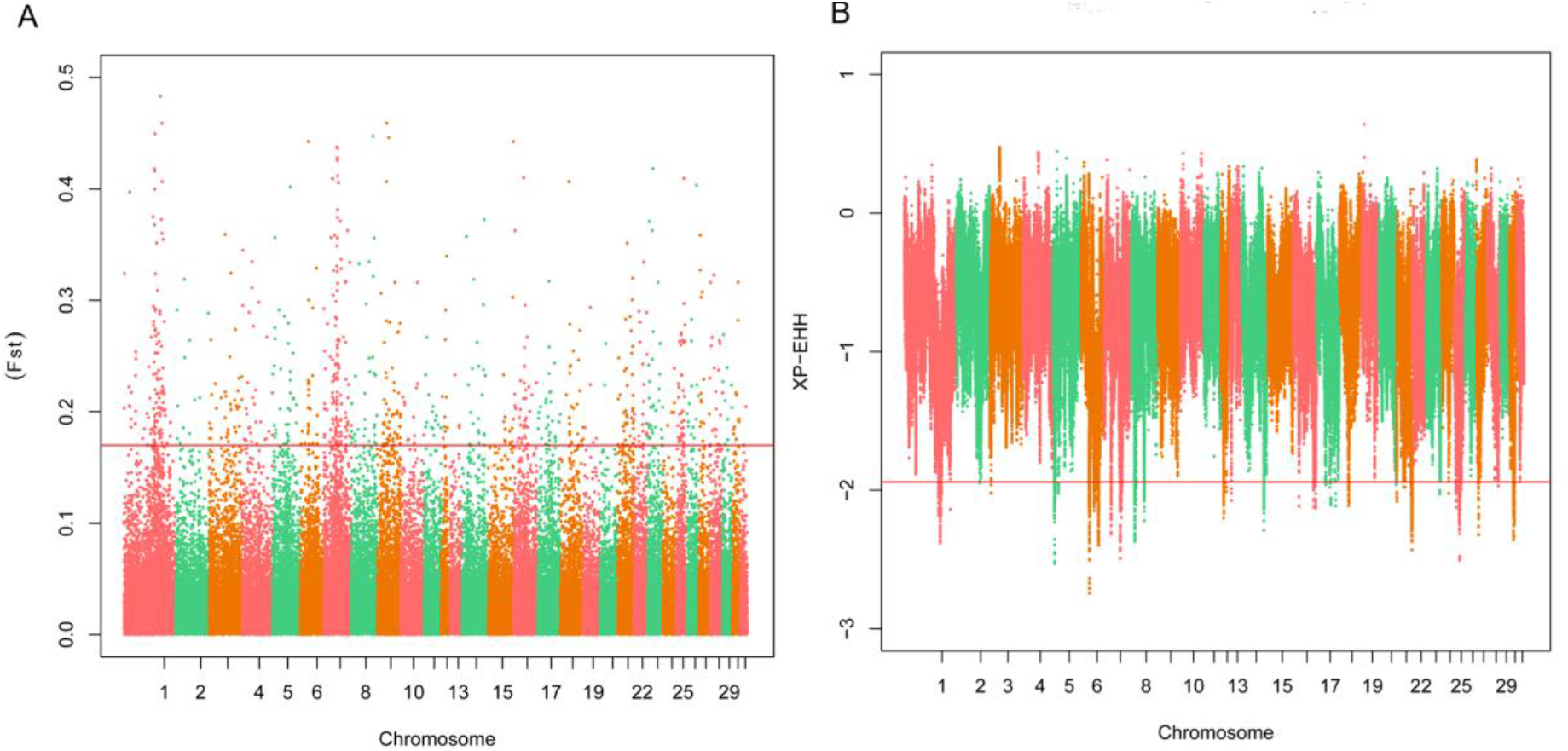
Selection signals of two population. (A) Genome-wide distribution of *F*_ST_ of RUL-YL on autosomes. Note: the scale on the X-axis represents ID of chromosomes, the scale on the Y-axis is the *F*_ST_ value of each SNPs. The red solid line indicates value of top 0.5%. (B) Genome-wide distribution of XP-EHH of RUL-YL on autosomes. Note: the scale on the X-axis represents ID of chromosomes, the scale on the Y-axis is the XP-EHH value of each SNPs. The red solid line indicates value of top 0.5%.

### XP-EHH test

The XP-EHH value (XP-EHH =-1.9405) was selected as the threshold. The SNP of largest XP-EHH value (AX-105009817, XP-EHH =-2.745) was located on chromosome 6. A total of 1,847 SNPs was selected as candidate sites, and the Manhattan plot was draw in Figure 2.

### Overlap analysis and gene annotation

A total of 34 SNPs was detected by both methods (Figure 3), 11 overlapped regions containing 56 genes were annotated (Table 1).

**Figure 3.**
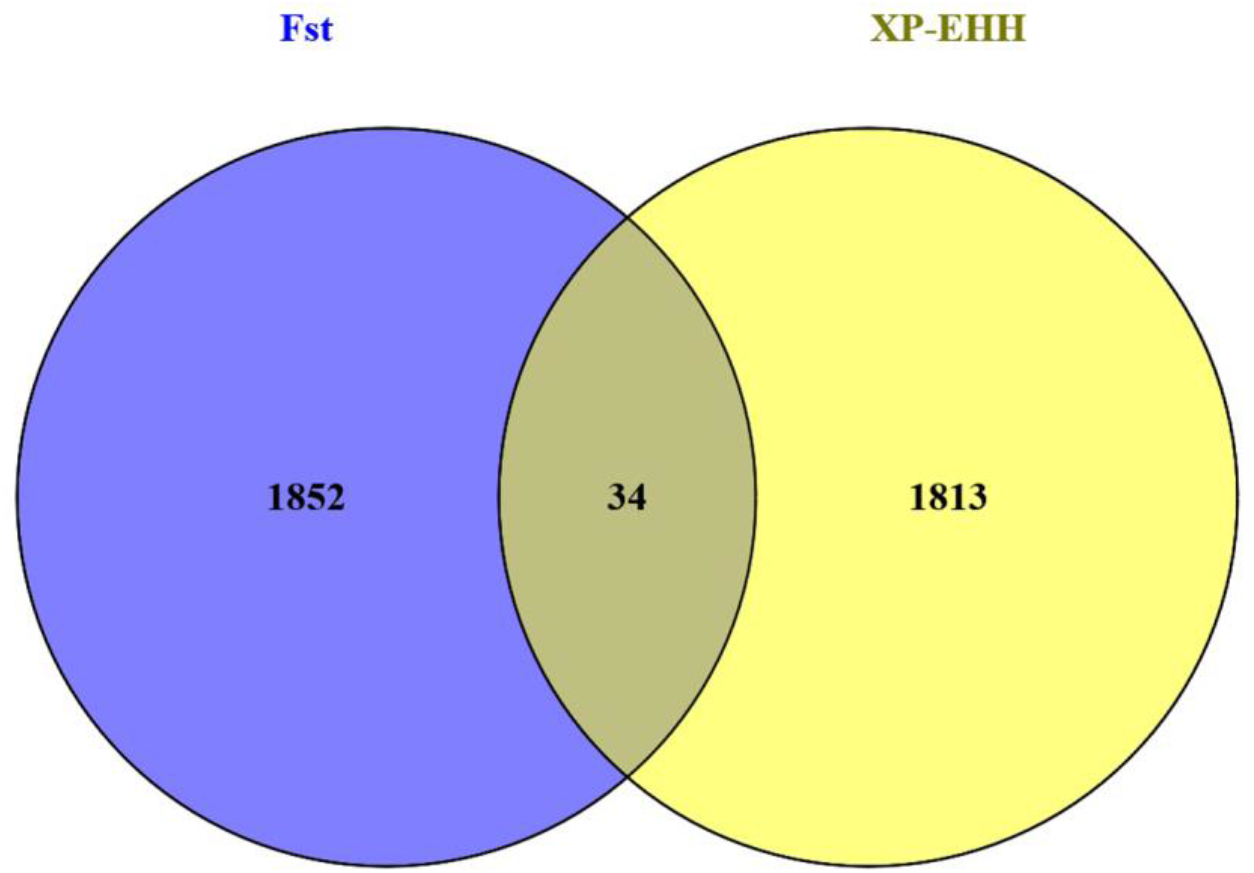
Venn diagram of two methods

**Table 1.**
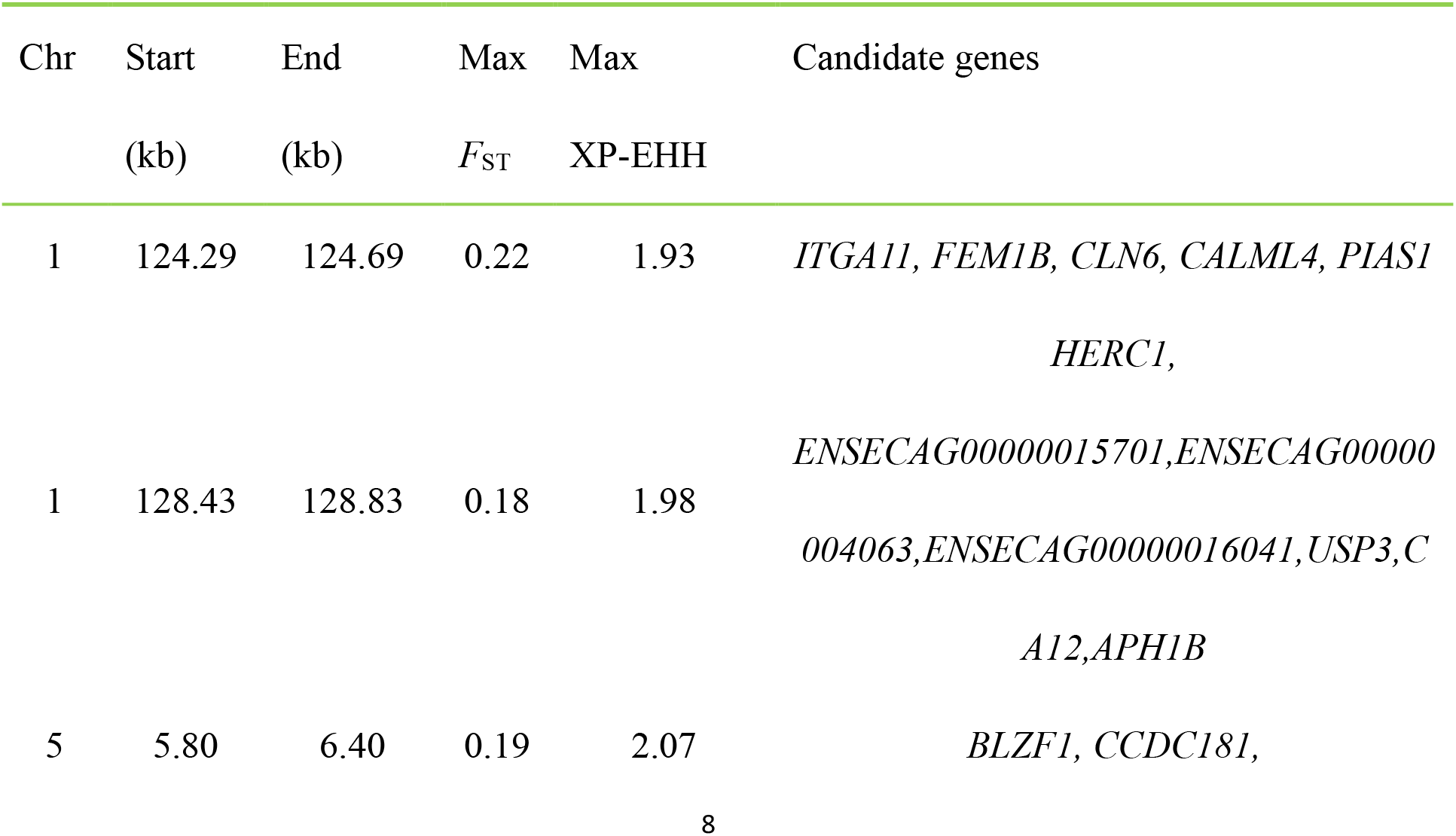
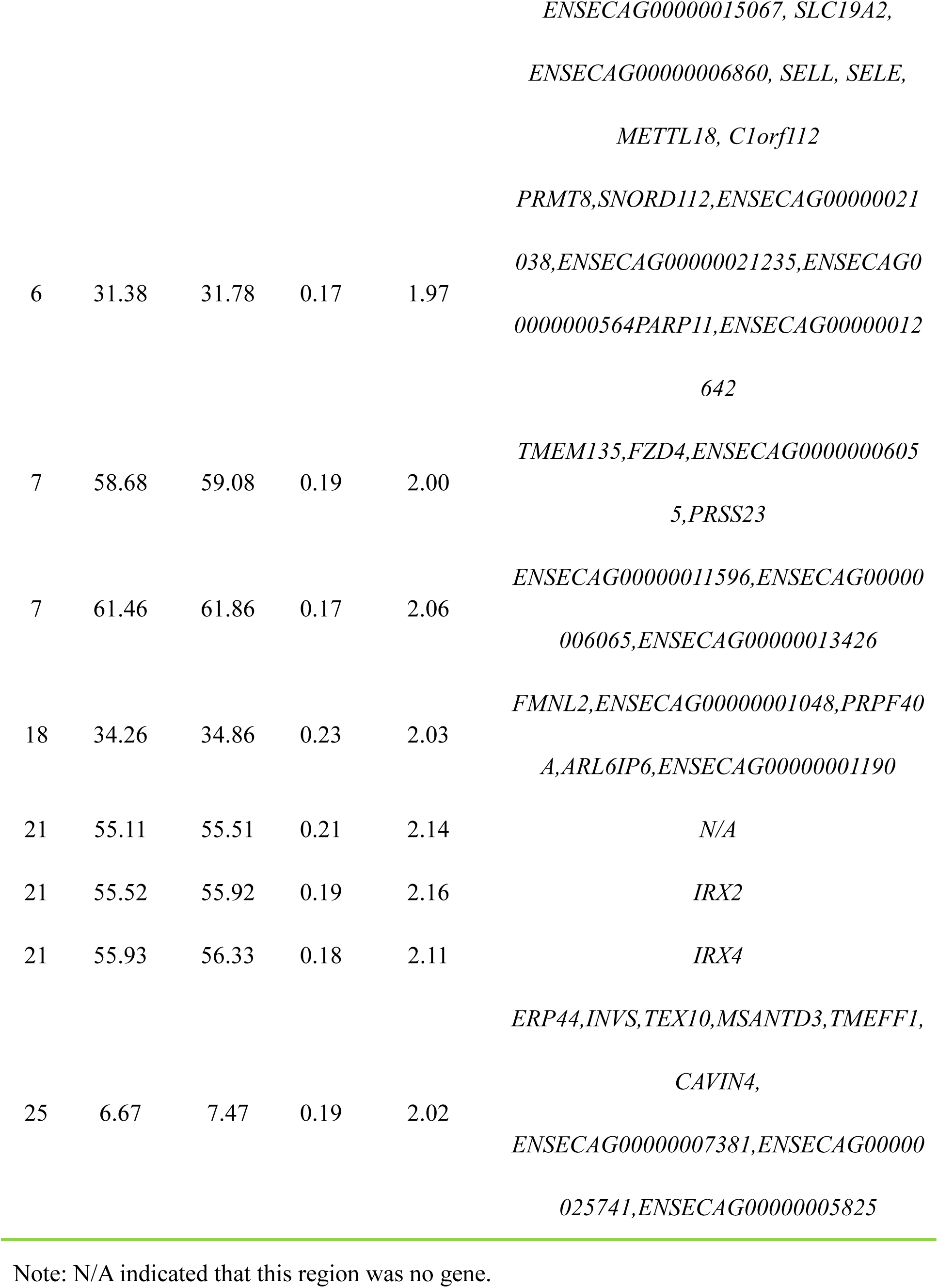
Selection regions and candidate genes of Yili horse population

### Gene enrichment analysis

DAVID 6.8 was used to enrich selected genes. Finally, we identified five significantly enriched categories (Table 2), and one term was may be related to gait of Yili horse.

**Table 2.** Significantly enriched annotation clusters and functional terms

| Software | Term | Category | Gene<br>count | P-value | Benjamini |
| --- | --- | --- | --- | --- | --- |
| DAVID | Selectin<br>superfamily | INTERPRO | 3 | 6.1e-6 | 4.9e-4 |
|  | Domain:EGF-like | UP-SEQ-FEATURE | 5 | 6.8e-6 | 1.1e-3 |
|  | C-type lectin,<br>conserved site | INTERPRO | 3 | 2.5e-4 | 2.5e-3 |
|  | Locomotion<br>involved in<br>locomotory<br>behavior | GOTERM_BP_DIRECT | 2 | 9.3E-3 | 4.6E-2 |
|  | oligosaccharide<br>binding | GOTERM_MF_DIRECT | 2 | 9.2E-3 | 3.8E-2 |

## Discussion

### Gait and Locomotion Performance

Gait of horse is the pattern of movement of the limbs, it can be divided into Natural gaits and Ambling gaits according to the formation reasons. Natural gaits are defined as the innate and untaught gait, such as walk, trot, pace, canter, gallop and jump. Ambling gaits are obtained by training, such as fox trot, paso fino, racking, running walk, tölt and slow gaits (Ensminger 1969). Gaits are often characteristic of a particular breed (for a kinematic study see Nicodemus & Clayton 2003) (Nicodemus *et al*. 2003), some horses are not able to learn a desired gait, others require extensive training, whereas some have a natural talent for ambling gaits.

Only DMRT3 gene is reported with horse gait. Anderson et al first show that a premature stop codon in the DMRT3 gene has a major effect on the pattern of locomotion in horses and find that Tölt is a unique ambling gait characteristic in Icelandic horse (Andersson *et al*. 2012). Subsequently, DMRT3 gene mutation was found to affect the Locomotion performance of horses in the Brazilian Mangalarga Marchador breed (Fonseca *et al*. 2017), the French trotter breed (Ricard 2015), American Saddlebred horses (Regatieri *et al*. 2016), Finn horses (Jäderkvist *et al*. 2015), Morgan and American Curly horses (Jäderkvist *et al*. 2014).

### Study on gait of china horse

In china, there are lots of horse breeds, but studies on gait are rare. Only Han et al. (2015) tested genotype of DMRT3 gene from 14 Chinese local horse breeds, The DMRT3 nonsense mutation occurs mainly in Northwestern China, where horses having a natural talent for pacing. Yili horses show good performance in Locomotion, Yili horse inherited the good locomotion performance from its ancestor Kazakh horse. But its specific mechanism remains to be studied. At present, there are no reports on the study of the gait in Yili horse. Considering the influence of genetic background and natural conditions, there may be unique mechanisms in gait. In this study, we found other genes may influence gait.

### Gene function

In this study, one term was fund associated with locomotory behavior, and CLN6, FZD4 gene were fund in this term. We know that the doublesex and mab-3-related transcription factor 3 gene (DMRT3) encodes an important transcription factor involved in the coordination of the locomotor system controlling limb movement (Andersson *et al*. 2012), and that the CLN6 gene is one of eight genes related to the liposome of nerve cells, the CLN6 gene comprises seven exons and is predicted to encode a 311 amino-acid protein with seven hypothetical transmembrane domains (Gao *et al*. 2002; Wheeler *et al*. 2002). The CLN6 gene was recently identified, mutations in which cause one of the variant late infantile forms of NCL (vLINCL) (Teixeira *et al*. 2003). This finding demonstrates a role for CLN6 in gait.

FZD4 is known to be involved in the Wingless (WNT) signaling pathway, which plays an essential role in cardiac development (Gessert *et al*. 2010). But this gene was no report in gait. The results of this study need further verification.

## Conclusions

In this study, we found eleven selected region, five enriched categories were found, one of them was associated with gait of Yili horse. The results of the present study indicate that the gait may be controlled by two genes (CLN6, FZD4) in Yili horse breed.

## Acknowledgements

We thank Yue-Hui Ma and Lin Jiang at institute of animal science, Chinese academy of agricultural sciences, Beijing. At the same times, we thank Ming-Jun Liu and Wen-Rong Li at the Xinjiang academy of animal husbandry for assistance with analysis platform of biological information data. This project was supported by the National Natural Science Foundation of China [Grant No.31560620]; National Sci-Tec R&D project of China [Grant No.2012BAD44B01].

## Ethics approval and consent to participate

All experimental procedures involving animals were approved (animal protocol numbers:2017001) by the Animal Care and Use Committee of Xinjiang Agricultural University, Urumqi, Xinjiang, China.

